# Patterns and perceptions of cannabis use with physical activity

**DOI:** 10.1101/328732

**Authors:** Jonathon K. Lisano, Kristina T. Phillips, Jeremy D. Smith, Matthew J. Barnes, Laura K. Stewart

**Author notes:** Corresponding author (LKS).

## Abstract

**Background and Aims:** Past research has shown that cannabis use is common among adults in the U.S. In addition, physical activity (PA), such as exercise, is often a component of many American’s daily routines. Anecdotal information suggests that a subset of individuals use cannabis in conjunction with PA, but the evidence base is lacking. The purpose of this study was to assess the frequency, methods of ingestion, strain types, and timing (before, during or after) of cannabis use in combination with PA. We also sought to better understand the types of PA that cannabis is being used with and reasons why individuals may use cannabis with PA.

**Methods and Results:** A brief survey was developed and was administered online to community residents (N = 105) who reported use of cannabis with PA. Analysis of survey responses revealed that participants were using cannabis in combination with a wide range of physical activities. While cannabis use was reported before, during, and after PA, the majority of participants (92%) reported use of cannabis before PA. Most participants (77%) believed that the use of cannabis products with their PA had a positive effect on their performance. The strain of cannabis used with PA was dependent on timing of cannabis use before, during, or after PA. Although participants reported a range of reasons for using cannabis before, during, or after PA, pain management was the only reason reported across all time periods.

**Conclusions:** Findings from this study suggest that there is a sub-community of physically active individuals using cannabis with their PA, many who believe that cannabis use has a positive effect on their performance.

## Introduction

Cannabis products, also commonly referred to as marijuana, are derived from the flower, stems and leaves of the hemp plant. In 2016, almost 9% of the U.S. adult population reported cannabis use within the past month, making cannabis the most commonly used illicit drug in the U.S. (1). Beginning in 2012 States within the U.S. began to legalize the recreational use of cannabis products. Currently, cannabis products are recreationally available for legal consumption in Alaska, California, Colorado, Maine, Massachusetts, Nevada, Oregon, Vermont, Washington and Washington D.C.

Phytocannabinoids, the active components in cannabis, mimic the effects of the endogenous cannabinoids in the body (2). Delta-9-tetrahydrocannabinol (THC) and cannabidiol (CBD) are the two most abundant phytocannabinoids present in cannabis products, and have received the most attention from the scientific community. However, THC and CBD are just two of more than 100 known phytocannabinoids (3) and the effects of these compounds have yet to be fully elucidated. Products of the cannabis plant can further be described by their cultivar, or strain, and are often separated into two general categories: *Cannabis Indica* and *Cannabis Sativa* (4), with varying hybrids of the two strains. Among medical cannabis users, common reasons for the use of *Cannabis Indica* include pain management and as an aid in sedation and sleep, while *Cannabis Sativa* users often prefer this strain for its perceived induction of euphoria and energy enhancement (5). Although exploration of the medicinal and psychoactive effects of cannabis products is still in its infancy, interest related to cannabis use on physical activity (PA) is also emerging.

Within the U.S., just over half (51.7%) of adults over the age of 18 years met the federal PA guidelines of at least 150-minutes of moderate or 75-minutes of vigorous activity per week (6). Currently, although there is minimal research describing how and why the physically active general population is using cannabis with PA, there have been several studies investigating cannabis use and athletic populations. Among male and female Division 1 National Collegiate Athletic Association (NCAA) athletes surveyed about their personal use of cannabis, 36.8% reported use within the past year (7). Thirty-eight percent of the athletes reporting cannabis use within past year reported using on average once per month, with male athletes more likely to report use when compared to their female counterparts (7). A more recent study found that athletes are more likely to use cannabis if they are male, Caucasian, or using performance enhancing drugs (8). The increasing availability of cannabis products for recreational use, combined with the growing number of U.S. adults who are physically active, and reports of athletes using cannabis may have contributed to a growing interest in the use of cannabis in combination with PA. More than 40 years ago, both moderate and heavy cannabis users were found to be less active the day after heavy cannabis use, however, it was speculated that the findings may have been associated with social reasons rather than pharmacological effects (9).

New evidence is emerging that the euphoric effects experienced during exercise, also termed as “runner’s high,” may be the result of the actions of endogenous cannabinoid release during exercise rather than endorphins (10). The G-protein coupled cannabinoid 1 receptors (CB1) in the brain have been observed to be closely linked to opioid receptors, and the dopaminergic reward pathways suggesting endogenous cannabinoid release with PA could be a major reason why regular exercise is perceived as highly rewarding (11, 12). It is possible that using cannabis products high in CB1 agonists, such as THC (13), could increase associated pleasure/reward already observed with regular exercise and increase motivation to partake in PA. Conversely, delayed-onset muscle soreness (DOMS) is often associated with muscle damage resulting from acute inflammation from strenuous exercise (14). Pain associated with DOMS may even result in exercise avoidance (15). -New evidence suggests that cannabinoids like THC and CBD are associated with pain reduction (16) and may have anti-inflammatory effects through their repressive effects on immune tissue (17). As a result, the use of cannabis may be a tempting option to reduce exercise-induced pain and inflammation. Yet, there is little evidence in human populations on how cannabis use combined with PA affects motivation to partake in exercise as well as how cannabis use affects recovery from exercise.

The primary goal of this exploratory study is to describe cannabis use as it relates to PA. More specifically, this study examines the frequency, method and timing (before, during or after) of cannabis use in combination with PA. Secondary goals include characterizing cannabis use as it relates to modes of PA and strain use and the examination of characteristics (e.g., age, gender) associated with participants’ cannabis use in conjunction with PA. Finally, we aim to better understand reasons that participants use cannabis with PA.

## Methods

### Participants and Procedures

From October to December 2017, 140 adults between the ages of 18 to 66 years across the U.S. were surveyed about their cannabis use habits in combination with their PA. Recruitment of participants was conducted online through social media (e.g., Facebook and Snapchat) and through snowballing and flyers posted in the local area for convenience sampling. A link and/or QR code took participants to a Qualtrics survey titled Cannabis Use and PA Questionnaire (CUPAQ), which they completed online. Recruitment materials specifically sought participants who use cannabis and cannabis products in relation to their exercise and PA habits. Participation was anonymous and took approximately 10 minutes. No external incentive was given for survey completion. This study was approved by the Institutional Review Board at the University of Northern Colorado.

### Survey Design and Administration

Initial contact in Qualtrics provided participants with a brief overview of the purpose of the study, emphasized that participation would remain anonymous, and noted the time required for participation (10 minutes). Participants were asked to complete an informed consent and confirm their age (18 or older) and residence in the U.S. prior to continuing. The survey consisted of 39 questions which were divided into three main sections. The first section consisted of 9 questions designed to gather general cannabis use habits of participants (i.e., frequency and duration of use), as well as age, gender, minutes of PA completed each week, and U.S. state of current residence. Section 2 included 18 questions focused on participants’ cannabis use habits as it pertained to their PA (before, during, or after PA). The frequency of cannabis use associated with PA over the last year and most recent episode of use were also assessed. Skip logic was programmed into Qualtrics so that participants who did not report cannabis use at one or more of the PA time points (before, during, or after) did not receive those questions. When cannabis was used before, during, and/or after PA, participants were asked to select the most common method of ingestion (e.g., smoking using a joint, inhaling via a vaporizer) and the strain (i.e., indica, sativa, or indica/sativa hybrid) if known. Participants were also asked to indicate the specific activities (e.g., weight lifting, kayaking) where cannabis was used before, during, and after PA. Lastly, an open-ended question assessed reasons for using cannabis before, during, and after PA. The third and final section of the survey consisted of 12 questions aimed at describing the amount and percentage of THC and CBD consumed. Using skip logic, questions were further divided into three categories based on participant self-reported primary form of cannabis use, including flower or bud, concentrates (i.e., oils, wax, shatter, dabs), and edibles. In assessing the quantity of the flower or bud, a visual aid and terminology were adapted from the Daily Sessions, Frequency, Age of Onset, and Quantity of Cannabis Use Inventory (DFAQ-CU) (18).

### Statistical Analysis

A total of 140 survey responses were obtained at the conclusion of the study. Three participants failed to complete informed consent and 32 participants reported never using cannabis in combination with their physical activities. These participants were removed from the dataset and a total of 105 participants were included in data analyses. All analyses were conducted using SPSS version 24 (IBM Corp.; Armonk, NY) and data are reported as frequency, percent or mean ± standard deviation. We present descriptive statistics to summarize the background characteristics of the sample and to examine participants’ use of cannabis before, during or after PA. To assess whether a range of demographic characteristics (e.g., age, gender) were associated with participants’ cannabis use during PA, we used a series of chi square analyses. Significance was p ≤ 0.05. Lastly, open-ended questions related to the reasons for using cannabis before, during, or after PA were examined through a content-analysis which allowed for the categorization of responses from each question into six or seven different themes. Responses for each of the three questions were coded independently by two coders into each of the theme categories. Agreement on these classifications was reached prior to listing a response under a specific theme category (prior to agreement, interrater reliability [k] = .80 −1.00). The frequency of responses was calculated and reported for each theme.

## Results

### Background Characteristics and Cannabis Use

Participants (53% male) ranged from 18-66 years of age (M = 31.4 ± 11.2 years) and lived in a total of 21 states across the U.S. The majority of participants were from Colorado (n=59), California (n=9), Louisiana (n=7) and Nebraska (n=5). Participants reported an average of 74.5 ± 111.5 months in duration of regular cannabis use. Ongoing cannabis use frequency revealed that 1.9% used less than once per month, 6.7% reported using between 1-3 times per month, 22.9% reported using between 1-6 times per week, and 68.6% reported using cannabis products on a daily basis.

### Physical Activity and Cannabis Use

Survey participants reported engaging in an average of 399.87 ± 543.82 minutes of PA throughout a typical week. The average age of participants when they first reported using cannabis with PA was 23 ± 8 years. When asked how frequently they used cannabis products in combination with PA, 63.8% of the participants reported using cannabis products in combination with PA within the past week. When asked to report their frequency of cannabis use in combination with PA, 9.5% used cannabis in combination with PA less than once a month, 12.4% used between 1-3 times per month, 41.0% reported using cannabis 1-6 times per week with PA, and 37.2% reported using at least one time or more per day in combination with their PA. Some participants who reported using cannabis with PA multiple times per day used before and after exercise, or used before exercise on two separate exercise bouts within the same day. The majority of participants (78.1%) reported using cannabis products in combination with PA at least once per week.

Participants also reported the method and quantity of cannabis used most frequently with PA. Methods/forms of cannabis consumption were grouped into four general categories: inhalation of flower/bud, edible, concentrate (dabbing), and other. The majority (80.0%) of participants reported that their primary method of cannabis use with PA was by inhalation of flower/bud. Methods of inhalation from most to least common are as follows: hand pipe (n=23), vaporizer (n=22), bong (n=18), joint (n=11) and blunt (n=6). Only 11.4% of participants reported primarily using concentrates with PA, 5.7% used edibles, and 2.9% other (topical/salves, capsules and fresh non-decarboxylated). Flower/bud forms were most frequently (n=80) reported when cannabis was used in conjunction with PA. Using the visual aid from the DFAQ-CU, participants who reported flower/bud as their primary form of use with PA personally used an average of 0.44 ± 0.45 grams of flower/bud before PA, 0.54 ± 0.49 grams of flower/bud during PA, and 0.78 ± 0.86 grams of flower/bud after PA.

### Timing of Cannabis Use with PA

When asked when they had used cannabis in conjunction with PA, 92% (n=97) of participants reported having used cannabis within one hour before beginning PA, 21% (n=22) reported having used cannabis during their PA, and 73% (n=77) reported having used cannabis within one hour after PA. Fifty-three percent (n=56) reported using most often before PA, 4.8% (n=5) reported using most often during PA, and 41.9% (n=44) reported using most frequently within one hour after completing their PA. A total of 23.8% of participants reported using cannabis only before PA, while 48.6% reported using before and after PA, followed by 18.1% reporting using before, during and after. Only 6.7%, 1.9% and 1.0% reported using only after, before and during, and only during respectively. The frequency of participant primary method of cannabis use before, during and after PA is presented in Table 1. Over three-fourths of participants who used cannabis either before, during, or after before PA reported their primary method of use was through inhalation. Smaller numbers used edibles or concentrates.

**Table 1.** Primary Method of Consumption Used Before, During and After Physical Activity

| Category | Method | Frequency Before<br>n (% of total) | Frequency During<br>n (% of total) | Frequency After<br>n (% of total) |
| --- | --- | --- | --- | --- |
| **Inhalation**<br>**(flower/bud)** | Joint | 12 (11.4%) | 15 (29.4) | 13 (13.8%) |
|  | Blunt | 7 (6.8%) | 4 (7.8%) | 7 (7.4%) |
|  | Hand pipe | 25 (24.3%) | 11 (21.6%) | 20 (21.3%) |
|  | Bong | 21 (20.4%) | 1 (2.0%) | 17 (18.1%) |
|  | Vaporizer | 20 (19.4%) | 12 (23.5%) | 14 (14.9%) |
| **Edible** | Edible | 5 (4.9%) | 3 (5.9%) | 2 (2.1%) |
| **Concentrate** | Dabbing | 11 (10.7%) | 4 (7.8%) | 14 (14.9) |
| **Other** | Other* | 2 (1.9%) | 1 (2.0%) | 7 (7.5%) |

Frequencies are reported in combination with the percentage of the total number of valid responses (Total (n)). \*Other forms of use included but were not limited to: topical/salves, capsules, and fresh non-decarboxylated.

The cannabis strain used most by participants before, during, and after PA is depicted in Table 2. Sativa and hybrid strains were most commonly used before PA, with hybrid and sativa strains most often used during PA, and indica and hybrid strains most commonly used after PA.

**Table 2.** Cannabis Strain Participants Had Used Before, During and After PA

| Strain | Before<br>n (% of total) | During<br>n (% of total) | After<br>n (% of total) |
| --- | --- | --- | --- |
| Indica | 20 (13.4%) | 8 (9.6%) | 57 (38.0%) |
| Sativa | 68 (45.6%) | 32 (38.6%) | 32 (21.3%) |
| Hybrid | 54 (36.2%) | 37 (44.6%) | 55 (36.7%) |
| Didn't Know | 7 (4.7%) | 6 (7.2%) | 6 (4.0%) |

Participants reported the strain(s) that they had used before, during and after PA.

### Perception of Cannabis Use on Athletic Performance

When participants were asked to report whether cannabis use with PA had a positive, negative, or no effect on their performance, 81 (77%) respondents reported they felt using cannabis in combination with their PA had a positive effect on their performance. A smaller number (n=21; 20%) reported that they felt cannabis use had no effect of their PA performance, and only 3 (3%) respondents felt cannabis use with their PA had a negative effect on their performance.

### Reported Physical Activities with Cannabis Use

Participants described using cannabis in association with both indoor and outdoor activities, as well as team and individual PA. Participants reported using cannabis before (Figure 1a), during (Figure 1b) and after (Figure 1c) a variety of PA. When participants used cannabis within 1 hour before PA, hiking (n=69), running (n=54), yoga (n=47), cycling (n=46), and resistance training (n=44) were the most commonly reported. The most frequent activities reported where cannabis was used during PA were: hiking (n=38), golf (n=19), yoga (n=16) and skiing/snowboarding (n=16). The most popular activities that participants reported using cannabis within 1-hour after completion of the activity were: hiking (n=51), running (n=49), resistance training (n=47) and cycling (n=39).

**Figure 1.**
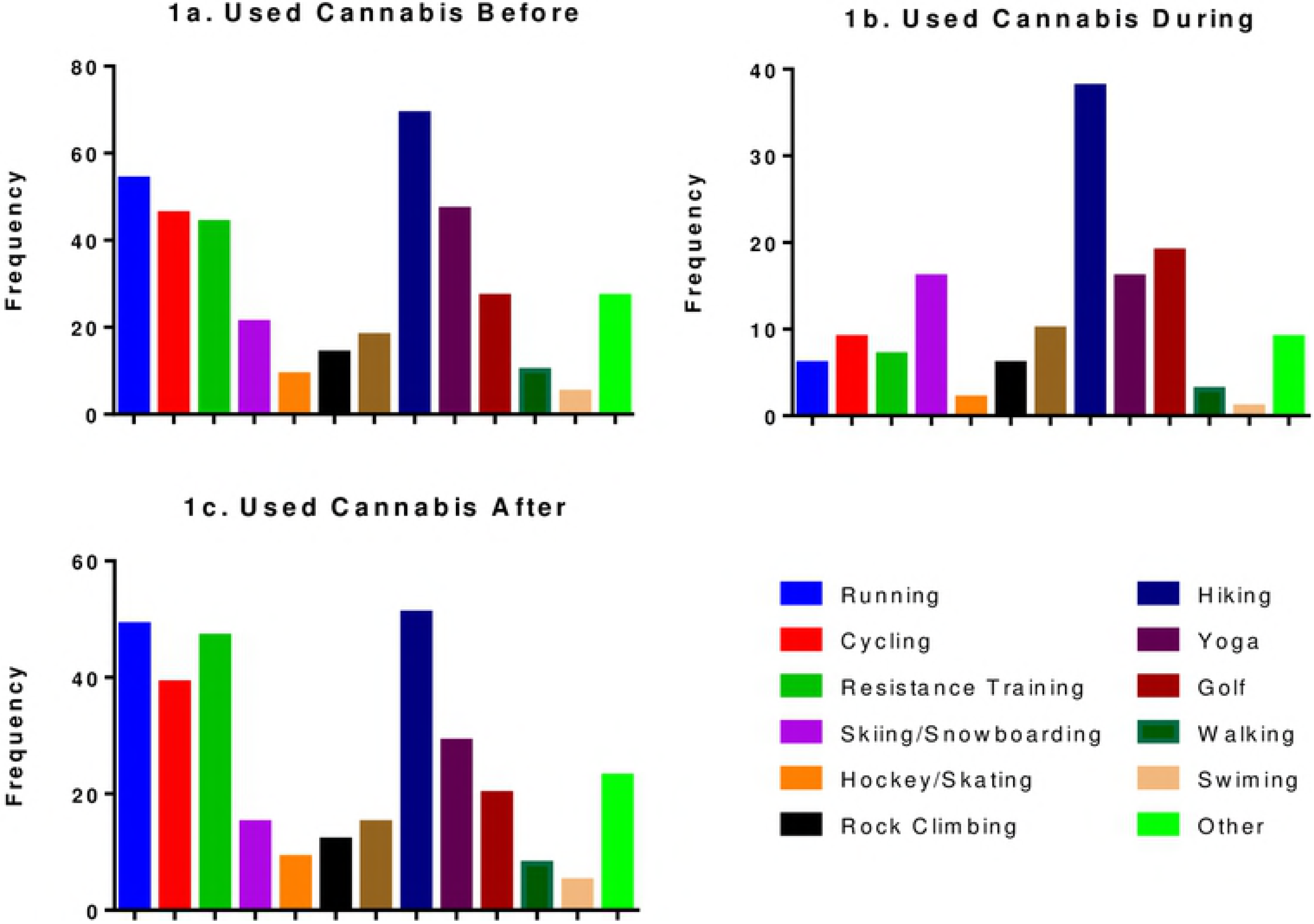
Frequency of Cannabis Use Before (1a.), During (1b.), and Alter (1c.) Physical Activities

### Demographic Characteristics associated with Marijuana Use and Performance

Chi-Square analysis revealed a significant difference between younger (≤ 27 years of age; n = 53) and older (≥ 28 years of age; n = 52) cannabis users with respect to their primary method of cannabis use with PA (p = 0.02). Older users favored more traditional methods of consumption via inhalation (i.e. joint, bong, pipe, vaporizer, and blunt), while younger users were more likely to use concentrates (dabbing). Younger users started using cannabis with PA at an earlier age (19.3 ± 2.9 years) when compared to older users (26.5 ± 10.5 years; p = 0.02). With respect to perceptions of cannabis use on performance, older users were more likely to report feeling that cannabis use had a positive effect on their PA performance. In contrast, younger users were more likely to report feeling that cannabis use had no effect on performance (p = 0.03). There were no significant differences between younger and older users with respect to the timing of cannabis use with PA (before, during, after; p = 0.44) or the frequency of cannabis use with PA (p = 0.74).

### Participant Reasons for using Cannabis with PA

Three separate open-ended questions asked participants to describe the reasons for using cannabis before, during, or after PA. Coded responses can be found in Table 3. Only participants who reported cannabis use during one or more of these times were provided with the respective open-ended questions. The three most common reasons for using cannabis before PA were: pain relief (n=25), to improve focus or get in the zone (n=25) and to calm the mind and/or body or to relax (n=25). The most popular reasons for using cannabis during PA was to increase/restore energy, push harder, or use as a break (n=12). Participants also described using cannabis during PA to improve their enjoyment of an activity and for pain management/relief (n=11). The overwhelming majority of responses for reasons of cannabis use after PA were related to relaxation (n=52) and to decrease pain or soreness (n=25), with minor reasons including appetite stimulation, and aid in sleep and recovery (Table 3). The only category that was present as a reason for cannabis use in all three times (before, during and after PA) was pain relief/management.

**Table 3.** Categorized Reasons of Cannabis Use Before, During and After PA

| **Reasons for Using Cannabis Before PA** | **Frequency (n)** |
| --- | --- |
| Pain management/relief | 25 |
| Improve focus, get in the flow, or “get in the zone” | 25 |
| Improve enjoyment of activity | 19 |
| Enhance performance, decrease fatigue, or to push harder | 16 |
| Improve motivation and state of mind | 18 |
| Calm mind and body; relaxation | 25 |
| Other | 18 |
| **Reasons for Using Cannabis During PA** | **Frequency (n)** |
| Improve enjoyment of activity | 11 |
| Improve focus, get back in or stay in the zone | 7 |
| Pain management/relief | 11 |
| Increase/restore energy, push harder, or to use during a break | 12 |
| Maintain the high | 4 |
| Other | 11 |
| **Reasons for Using Cannabis After PA** | **Frequency (n)** |
| Relaxation | 52 |
| Stimulate/increase appetite | 7 |
| Pain management/relief | 25 |
| Aid in sleep | 6 |
| Aid in recovery | 9 |
| Other | 18 |

## Discussion

The present study aimed to describe how and why physically active individuals are using cannabis in combination with PA. Not only are individuals using cannabis in combination with PA, but individuals are using cannabis products before (within 1-hour of starting PA), during, and after PA (within 1-hour of cessation of PA). Studies assessing cannabis use in elite athletes have found that individuals who were male, Caucasian, and played hockey were most likely to report cannabis use (7, 8). However, there is little data exploring the characteristics of cannabis use in recreationally physically active individuals. The present study found that cannabis use with PA was reported equally among males and females and across a wide range of activities in recreationally physically active individuals. With a self-reported range of 25-3600 minutes of PA per week, 73 of the 105 participants met the national weekly physically activity requirements. Hiking was reported as the most prominent activity to use cannabis with before, during, and after PA. Further observation suggests that the timing of cannabis use is heavily dependent on the specific PA. Running, cycling, and resistance training were the 3^rd^, 4^th^, and 5^th^ most frequently reported with cannabis use prior to activity, and were the 2^nd^, 3^rd^, and 4^th^ most frequently reported activities to use cannabis after. However, reported frequency of cannabis use during running, cycling, and resistance training dropped to 6^th^, 7^th^ and 8^th^ respectively. The most popular reported activities to use cannabis during PA were: hiking, golf, skiing/snowboarding, and yoga. Together, these results suggest that the timing of cannabis use with physical activity is heavily dependent on mode of PA.

The majority of participants were under the impression that cannabis use in combination with PA had a positive effect on their PA performance, while only 3% felt that using cannabis had a negative effect on PA performance. The perception of improved performance with cannabis use may be purely subjective. The most recent studies which examined this question were conducted 40 years ago and demonstrated that acute use of cannabis containing THC increased resting heart rate (19, 20), systolic and diastolic blood pressure (20) as well as reduced time to exhaustion during a cycling bout (21). Yet, no acute effects of THC administration were reported with respect to oxygen uptake or ventilation during submaximal exercise (19). Unfortunately, the concentration of THC in the cannabis used in these studies is no longer reflective of current cannabis products available on the market today, as THC content in cannabis has been steadily increasing over the past several decades (22).

While evidence is lacking related to assessing the effects of acute consumption of cannabis on exercise performance, a recent study explored the effects of chronic cannabis use on exercise performance. In this study, participants were assessed for pulmonary, cardiovascular, anaerobic and strength while at least 12-hours removed from last use of cannabis. When compared to a non-cannabis using control group this cross-sectional study found that there were no differences with respect to pulmonary, cardiovascular, anaerobic, or strength performance (23). However, findings from the present study revealed that a large portion of participants believed that cannabis use had a positive effect on their performance. More research is needed to fully elucidate if the acute use of cannabis is actually effects performance.

Previous research has shown that adults aged 18-25 have the highest reported percentage of cannabis use (1), with 20.8% reporting use at least once within the past month (1). While the current study did not assess cannabis use rates among age groups of physically active adults, an age-based analysis was done to assess if individuals 18-27 years of age had different perceptions and methods of use compared to adults over 28 years of age. Chi-square analysis revealed that adults ages 18-27 were significantly more likely to report using concentrates as their primary form of cannabis and began using cannabis with PA at a significantly younger age (i.e., 7 years earlier). However, adults over the age of 28 were significantly more likely to report feeling that cannabis use had a positive effect on their performance compared to adults aged 18-27. Emerging research shows that use of concentrates via dabbing may be associated with greater negative consequences, tolerance, and withdrawal compared to flower use (24). We did not assess negative consequences related to participants’ cannabis use, but future research should explore whether those using cannabis with their PA are more likely to have problems related to their use and if they are more likely to use cannabis with PA for specific reasons (e.g., pain management).

Results from this study revealed that 92% of survey participants reported cannabis use within 1-hour of beginning PA, suggesting that more research to ascertain the effects of acute cannabis use on PA performance may be necessary. Conversely, the perceived performance enhancing effect of cannabis on PA performance could be related to the reduced perception of pain. When asked an open-ended question as to why participants used cannabis before, during, and after PA, pain management/relief was the only reason to be reported across all time points. Pain management was the most common reason for cannabis use before PA, and the second most common reason for use during and after PA. This pain control theme is supported by a recent study which found that pain was the most commonly reported reason for seeking use among medicinal cannabis users (25, 26) with those seeking pain relief preferring *Cannabis Indica* (5, 27). Products derived from *Cannabis Indica* are typically lower in THC and container higher quantities of CBD, reducing the perceived psychoactive effects while still maintaining high pain suppressive effects. Mechanistically cannabinoids modify synaptic transduction in the central nervous system and the periphery. THC and CBD are agonists of the two primary cannabinoid receptors, CB1 and CB2 (2), with CB1 being highly expressed in the central nervous system (28) and CB2 more abundant in the periphery (29). These cannabinoids act on CB1 and CB2 receptors expressed on the pre-and post-synaptic membrane blocking calcium influx, and blocking synaptic vesicle release (30). Activation of these receptors blocks synaptic signal transduction and has even been implicated in long-term depotentiation (31). This mechanism provides a suitable explanation as to why cannabis is used for exercise-associated pain reduction. Interestingly, individuals using cannabis for pain mediation have been found to be at lower risk of development of cannabis use disorders (27).

In conjunction with pain management, the most common reasons reported for cannabis use prior to PA were related to improved focus and to calm or relax the mind and body. This was unexpected, as *Cannabis Sativa* was the most frequently reported strain used prior to PA, which is typically associated with feelings of euphoria and energy enhancement (5). After PA, using cannabis for relaxation was reported more frequently than any other response. Given the reported perceived effects of *Cannabis Indica* related to sedation and pain management (5), it was expected that this strain may be used predominately post-exercise. This was supported in this study with only 13% and 10% of participants reporting the use of *Cannabis Indica* before and during exercise, respectively. *Cannabis Indica* predominant strains were the most frequently used strain following PA at 38%. This practice could be influenced by perceptions of the effects of *Cannabis Indica* that are popular in the cannabis community.

The method and form of cannabis appears to differ based on the time when cannabis is used in relation to PA. Most participants reported consuming cannabis through traditional inhalation methods regardless of PA time point. Yet, the distribution of individual inhalation methods at each of those time points did vary. Before PA, cannabis consumed with a hand pipe was the primary method of use by nearly a quarter of participants. Comparatively, 30% reported using a joint and 23% described using a vaporizer during PA. Primary flower/bud inhalation methods then shifted back to predominant hand pipe and bong use following completion of PA.

Although this study provides new information on why physically active individuals are using cannabis products with their PA, it does have a number of limitations. The study was cross-sectional and conducted as an online survey. There are limitations associated with self-report data, even though online and in-person administration of surveys has been shown to yield similar results (32). Reported weekly minutes of PA varied dramatically, with a range of 25 to 3600 minutes of physical activity per week. In an effort to allow participants to report any type of physical activity, participants were not asked if they were physically active for recreational reasons, or if their PA was a part of a structured exercise regimen. Although this approach allows for more broad interpretation of physical activity, it should be considered a limitation and future work should further examine this question. In addition, the present study did not explore if participants experienced any negative side effects due to their cannabis use, such as those associated with Cannabis Use Disorder. Future studies exploring the use of cannabis with exercise may want to discern whether individuals are consuming *ad libitum* and happen to be physically active vs. those who are intentionally using cannabis in conjunction with structured exercise.

In summary, this study provides novel insight into cannabis use among individuals that reported using cannabis in combination with their physical activity. This study revealed that the most common time to use cannabis in combination with PA was within 1-hour of starting PA, with the majority of individuals reporting use through traditional inhalation methods. The majority of participants reported feeling that the use of cannabis with PA had a positive effect on PA performance. Reasons for cannabis use with PA were heavily dependent on timing of cannabis use in relation to that activity with pain management as the only common reason reported at before, during, and after time points. Finally, results from this study indicate that timing, method of use and strain of cannabis use were commonly considered when cannabis was used in conjunction with PA.

## Acknowledgements

The authors would like to thank the University of Northern Colorado for the support of this project, and for the use of the Qualtrics software. We would also like to thank our participants for volunteering their time to participate in this study. Finally, we would like to thank the reviewers for their helpful comments in the revision of this work.

